# Towards implementation of AI in New Zealand national screening program: Cloud-based, Robust, and Bespoke

**DOI:** 10.1101/823260

**Authors:** Li Xie, Song Yang, David Squirrell, Ehsan Vaghefi

## Abstract

Convolutional Neural Networks (CNN)s have become a prominent method of AI implementation in medical classification tasks. Grading Diabetic Retinopathy (DR) has been at the forefront of the development of AI for ophthalmology. However, major obstacles remain in the generalization of these CNN’s onto real-world DR screening programs. We believe these difficulties are due to use of 1) small training datasets (<5,000 images), 2) private and ‘curated’ repositories, 3) offline CNN implementation methods, while 4) relying on accuracy measured as area under the curve (AUC) as the sole measure of CNN performance.

To address these issues, the public EyePACS Kaggle Diabetic Retinopathy dataset was uploaded onto Microsoft Azure™ cloud platform. Two CNNs were trained as a “Quality Assurance”, and a “Classifier”. The “Classifier” CNN performance was then tested both on ‘un-curated’ as well as the ‘curated’ test set created by the “Quality Assessment” CNN. Finally, the sensitivity of the “Classifier” CNNs was boosted post-training using two post-training techniques.

Our “Classifier” CNN proved to be robust, as its performance was similar on ‘curated’ and ‘uncurated’ sets. The implementation of ‘cascading thresholds’ and ‘max margin’ techniques led to significant improvements in the “Classifier” CNN’s sensitivity, while also enhancing the specificity of other grades.

## Introduction

It is estimated that by 2040, nearly 600 million people will have diabetes worldwide(1). Diabetic retinopathy (DR) is a common diabetes-related microvascular complication, and is the leading cause of preventable blindness in people of working age worldwide(2, 3). It has been estimated that the overall prevalence of non-vision-threatening DR, vision-threatening DR and the blinding diabetic eye disease were 34·6%, 10·2%, and 6·8% respectively (3-6). Clinical trials have shown that the risk of DR progression can be significantly reduced by controlling major risk factors such as hyperglycaemia and hypertension (7-9). It is further estimated that screening, appropriate referral and treatment can reduce the vision loss from DR by 50% (10-12). However, DR screening programs are expensive to set up and administrate. It is estimated that even in developed countries, these programs do not reach up to 30% of the diabetic population (13, 14).

Artificial intelligence (AI) and its subcategory of deep learning have gained popularity in medical screening programs, including DR screening. In deep learning, a convolutional neural network (CNN) is designed and trained based on large datasets of ground truth data and labels. The CNN algorithm adjusts its weights and discovers which features to extract from medical data (e.g. fundus photos) to achieve the best classification accuracy, when compared to human performance (15-20). CNNs use layers with convolutions, which are defined as mathematical functions that use filters to extract features from an image (21-23). The output of a DR classifying CNN can be either a binary classification such as Healthy vs Diseased; or a multiclass classification task such as Healthy, Non-referable DR, Referable DR (16, 24).

The rapid initial advances of AI, especially in DR classification, have hyped the immediate implementation of AI in national DR screening programs, and subsequent noticeable cost savings (25, 26). However, these systems have not yet been successfully translated into clinical care, due to major generalizability issues of research-built AIs. Some of the major flaws of research-built AIs that are hindering their generalizability are 1) using small training (<5,000 images) datasets, 2) repositories that are often private and ‘curated’ to remove images that are deemed to be of low quality, and 3) lack of external validation (17, 27-29). These issues are often observed in research-driven AIs and have led to a slew of extremely biased DR classifying neural networks in the published literature. Some recent publications have pointed out the lack of generalizability of even the best of these AIs (30-33).

Our extensive investigation (to be published soon as a systematic review) have found only a few published research-based AIs that could be closer to clinical translation (4, 16, 18, 19, 26, 34-36). Although great works in their own right, these AIs often need dedicated and TensorFlow compatible graphic cards (GPUs) to achieve rapid live image grading. Often, public health providers rely on older and\or less expensive IT infrastructure, so such high computational demand would hinder their clinical translation. Such implementation is important since, to access more than 30% of the diabetic population are not reached (even in advanced countries with established screening programs), especially in remote, rural and low socioeconomic regions.

Finally, the creators of DR-screening AIs have traditionally focused on improving the accuracy of their trained AIs, as measured by the area under the curve (AUC)(17). Although reasonable, it should be noted that different diabetic eye screening programs will have different requirements. It could be argued that 1) the emphasise of a community-based screening program, potentially operating in the remote and low socioeconomic region and using portable handheld cameras, is on identifying those patients with no disease from those with any disease. However, in traditional screening, a CNN which is highly sensitive and removes the need for a significant (>70%) portion of images to be sent for human review, would lead to immediate and significant cost savings for the program.

We are actively working towards implementation of our DR classifying AI, within a long-established diabetic eye-screening program in New Zealand(37, 38). This program has multiple facets, including community-based and clinic-based screening phases. In this project, EyePACS Kaggle Diabetic Retinopathy public dataset was used to develop two CNNs, based on one of the most sophisticated architectures available. Next, both CNNs were deployed and trained on the Microsoft Azure™ cloud platform as 1) a fundus image “Quality Assessment” and 2) a DR “Classifier”. The “Quality Assessment” CNN was used to create a ‘curated’ test set, in addition to the original ‘un-curated’ set. The performance of the “Classifier” CNN was assessed on both sets. Finally, we used two post-training methods to boost the sensitivity of the “Classifier” CNN towards 1) Healthy grade and 2) the most severe DR. We are actively pursuing clinical implementation of our AIs and our recent findings would be of great interest for similar groups around the world.

## Methodology

The original EyePACS Kaggle DR was obtained and uploaded onto the Microsoft Azure™ platform. Initially, a “Quality Assessment” CNN was trained for assessing the quality of the retinal images. Next and to better match the New Zealand grading scheme, the original grading was modified in two ways [Healthy vs Diseased] and [Healthy vs Non-referable DR vs Referable DR]. The uploaded dataset was then divided into training (70%), validation (15%) and test (15%) sets. A separate “DR Classifier” CNN was then trained on the Microsoft Azure™ platform, using the ‘un-curated’ training and validation datasets. The not-seen-before test set was then analysed by the “Quality Assessment” CNN thus creating a ‘curated’ test set in addition to the original ‘un-curated’ set. The performance of the “Classifier” CNN was then assessed using both ‘curated’ and ‘un-curated’ test sets. Finally, the ‘cascading threshold’ and ‘margin max’ techniques were implemented post-training, to investigate their effects on boosting the sensitivity of the “Classifier” CNN [Figure 1].

**Figure 1:**
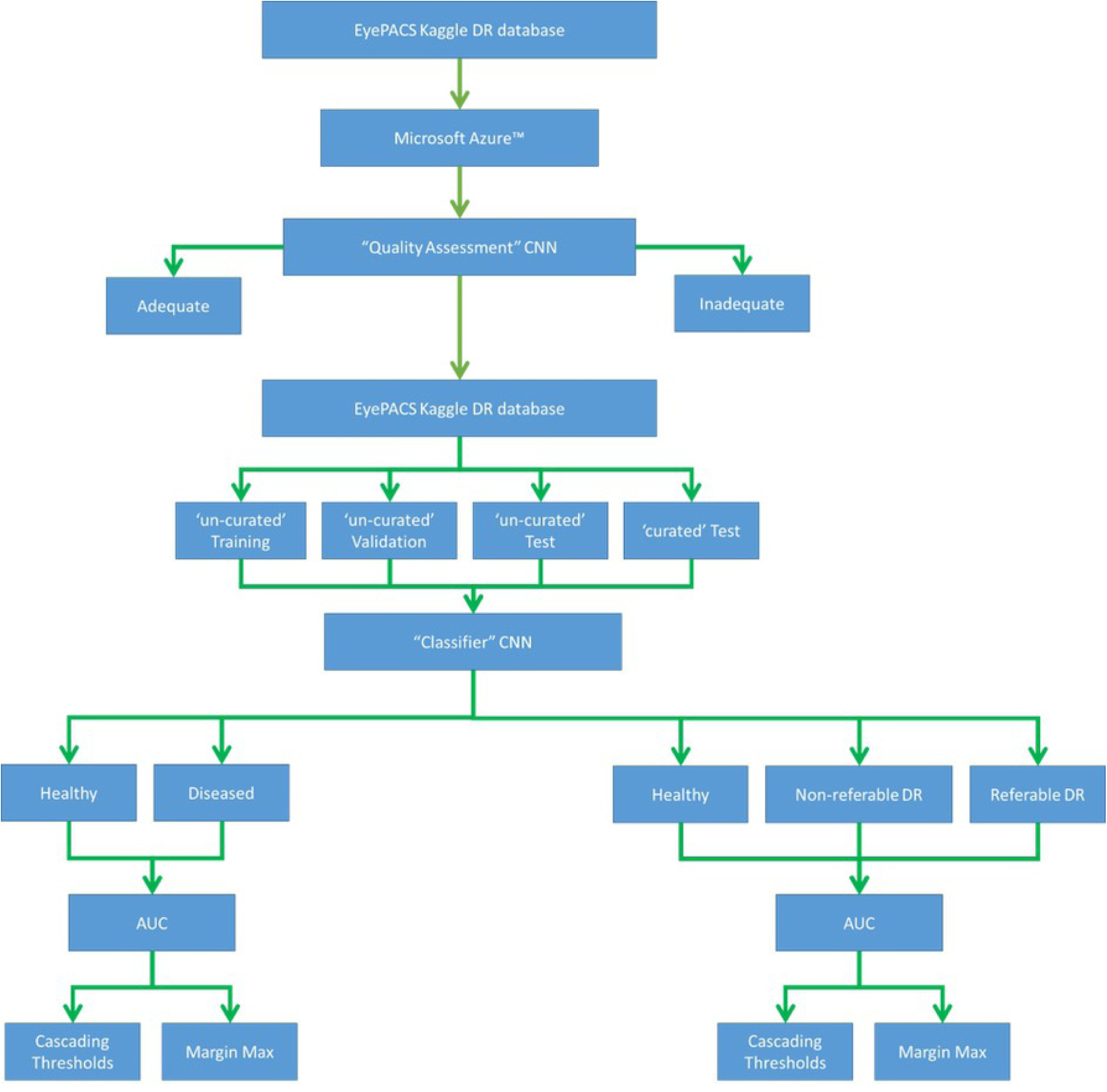
Flowchart of our AI design, implementation and test. The public data atteiment and upload onto the Microsoft Azure cloud platform was the first step. “Quality assessment” CNN was trained to identify adequate and inadequate images. the entire public dataset was then devided to training, validation and test sets. The test set was then ‘curated’ by the “Quality assessment” CNN. The “Classifier” CNN was trained on un-curated data, and then tested on ‘curated’ and ‘un-curated’ data. Its erformance was also assessed using 2 or 3 DR lables.

### Quality Assessment Dataset

A subset of 7,026 number of images from the original set were used for creating the “Quality Assessment” CNN. The images were audited by a senior retinal specialist (DS) and labelled as ‘adequate’ or ‘inadequate’ (3400 \ 3626) respectively [Figure 2]. They were then split into (75%) training, (15%) validation and (15%) testing sets.

**Figure 2:**
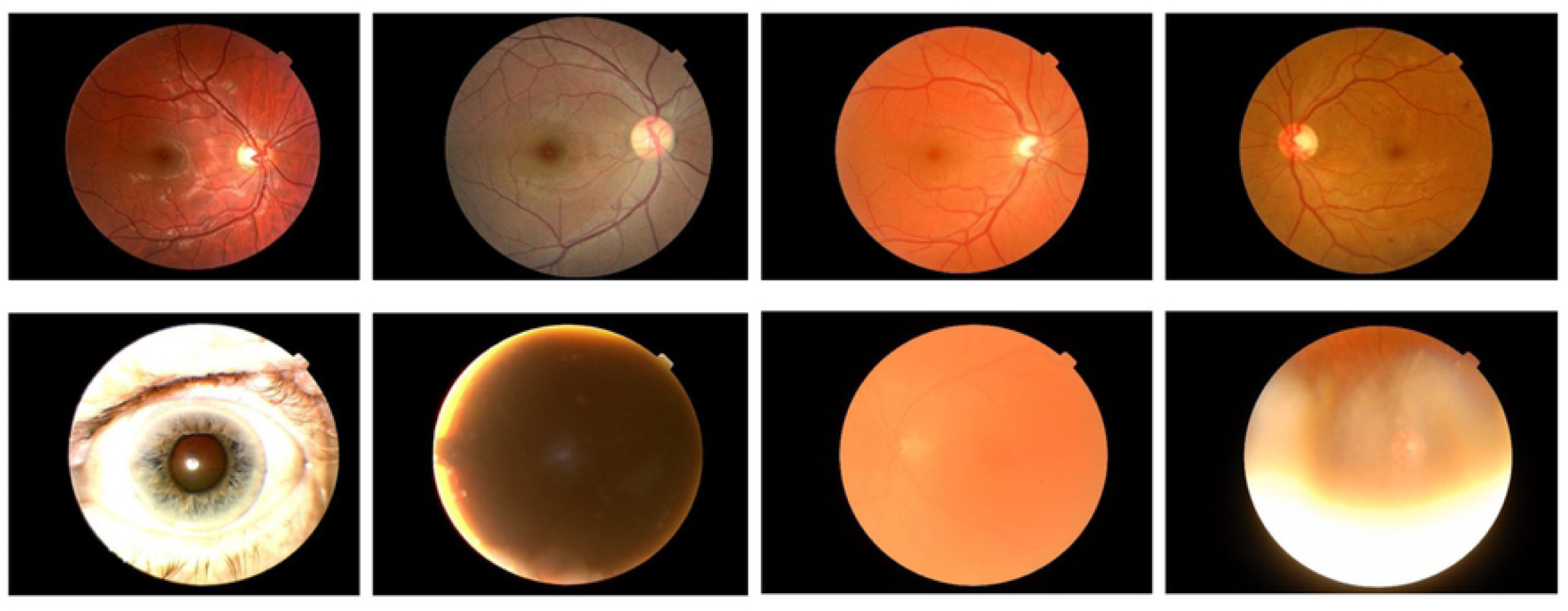
samples of ‘adequate’ and ‘in-adequate’ images as decided by a senior retinal specialist. Fundus images deemed adequate are shown in the upper row. Fundus images deemed inadequate are shown in the bottom row.

### “Quality-Assessment” CNN Architecture

To choose the optimum CNN design, several architectures were tested on Microsoft Azure™ cloud platform. These included ResNet, DenseNet, Inception and Inception-ResNet and Inception-ResNet-V2 (39). The “Quality-Assessment” CNN was then based on a modified version of the InceptionResNet-V2 architecture. For our purposes, the number of neurons in the final output layer was changed to two, corresponding to ‘adequate’ and ‘inadequate’ classes. The learning rate was 0.001, using ADAM optimizer, with a mini-batch size of 30, and training was continued to 100 epochs.

### Classifier Dataset

The public Kaggle Diabetic Retinopathy was downloaded through EyePACS, which can be found in https://www.kaggle.com/c/diabetic-retinopathy-detection/data. This dataset contains 88,700 high-resolution fundus images of the retina, labelled as No DR, Mild, Moderate, Severe, Proliferative DR. To mimic the decision making of the New Zealand national DR Screening program, the original grading was remapped to three cohorts of Healthy, Non-referable DR and Referable DR [Table 1]. Furthermore, as one potential gain of using an AI in DR Screening program is to quickly identify those that are healthy, separately the dataset was remapped to the broad classification of Healthy vs Diseased [Table 2].

**Table 1:** re-categorization of the original Kaggle EyePACS grading scheme (5 grades) to three new categories

| Original Grade | New Grade | Number of images |
| --- | --- | --- |
| No DR | Healthy | 65,300 |
| Mild DR | Non-referable DR | 19,400 |
| Moderate DR |  |  |
| Severe DR | Referable DR | 4,000 |
| Proliferative DR |  |  |

**Table 2:**
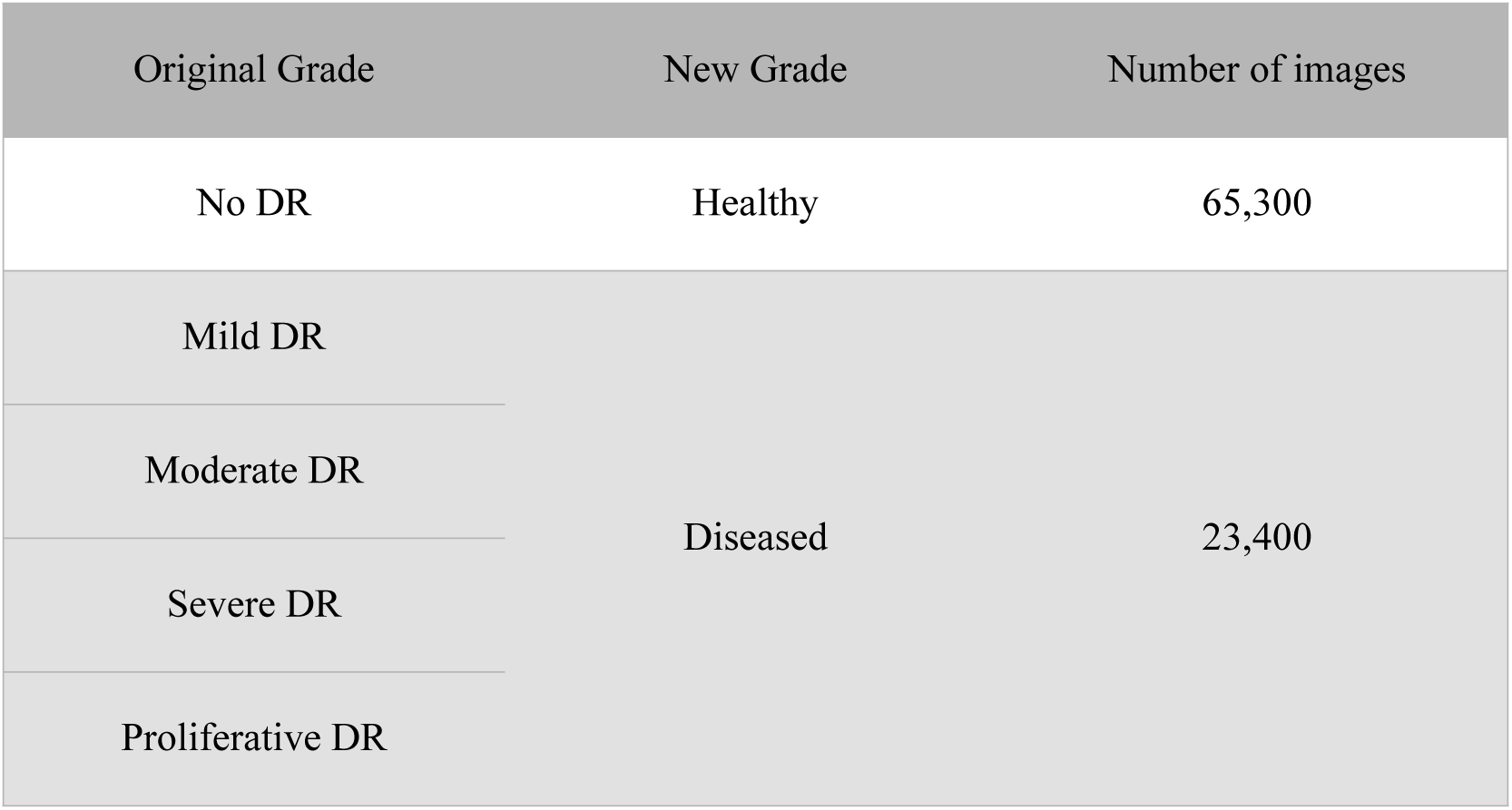
re-categorization of the original Kaggle EyePACS grading scheme (5 grades) to two new categories

| Original Grade | New Grade | Number of images |
| --- | --- | --- |
| No DR | Healthy | 65,300 |
| Mild DR | Diseased | 23,400 |
| Moderate DR |  |  |
| Severe DR |  |  |
| Proliferative DR |  |  |

Each re-categorized dataset was then split into training set, validation set and testing set with corresponding ratios of 70%, 15% and 15% respectively. The final number of images were 62,090 for un-curated training set, 13,305 for un-curated validation set and 13,305 and 6,900 for ‘un-curated’ and ‘curated’ test sets respectively.

### Pre-processing

The Kaggle EyePACS images were cropped and resized to 600*600. The choice of image size was to minimize the computational load on the Microsoft Azure™ platform, while not compromising the performance of the trained CNNs. According to existing literature (40) and based on our experience, larger image sizes would have led to diminishing returns in accuracy and overall performance of the designed CNNs. The resized cropped images were enhanced by applying a Gaussian blur technique (41), using the equation below.

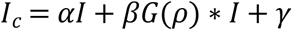

A series of Gaussian blur parameters were tried and an optimum set was chosen by a senior retinal specialist (DS):

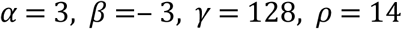

The Gaussian blur technique has been designed to remove the variation between images due to differing lighting conditions, camera resolution and image qualities [Figure 3].

**Figure 3:**
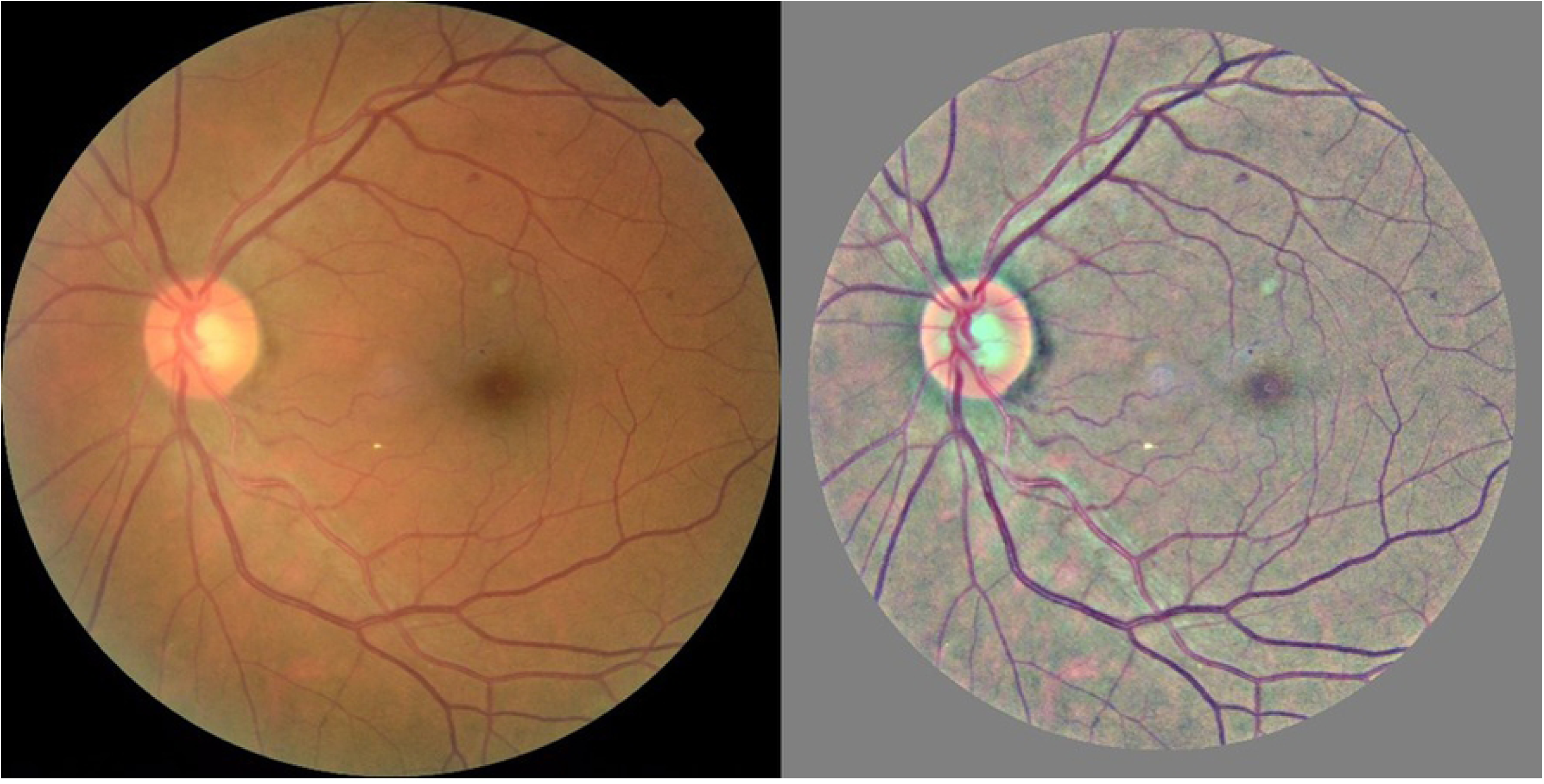
Contrast enhancement of the Kaggle EyePACS fundus image. The using Gaussian blur technique was applied to the raw fundus image (left). This technique minimizes intensity and contrast variability in fundus image dataset (right).

### “Classifier” CNN Architecture

The “Classifier” CNN was then designed based on the Inception-ResNet-V2 architecture, since this architecture has outstanding network capacity, faster convergence speed and better stability, which are critical when training utilizing such a large dataset. Three sequential layers of a GlobalAveragePooling2D layer, a dropout layer (dropout rate = 0.3) and a fully connected layer were added to the original architecture. The activation function of the added dense layer was a Softmax function; and cross-entropy loss/error was utilized as the loss function, while Adam algorithm was utilized as the optimizer. The learning rate was 0.001 and a mini-batch size of 64 was used for model training, and training was continued for 100 epochs. Finally, a weighted loss-function was used here to address the class imbalance of the Kaggle EyePACS dataset.

### Cascading Thresholds

The cascading thresholds technique has been used previously in the literature, in order to boost the sensitivity of a given CNN (33). Normally, a classifying CNN has a Softmax layer followed by a Classification Output layer as its last layers. The Softmax layer generates a list of probabilities of given input (i.e. fundus photo), to belong to a certain class (i.e. Healthy, Non-referable DR, Referable DR). The Classification Output layer will then choose the class with maximum probability as the outcome of the classifier CNN. Alternatively, to increase the sensitivity of the CNN towards a specific grade (e.g. Referable DR), sub-maximum probabilities of that specific grade could be used.

An example of (∼, 0.3, 0.3) cascading thresholds limit is presented here. Following AI’s image analysis, if the output of the Softmax layer for the Referable class reaches the threshold of 0.3, then regardless of the less severe grades probabilities, the image is classified as Referable. If the image is not classified as Referable and if the Softmax layer output of Non-referable DR grade reaches the threshold of 0.3, this image is then assigned to this grade, regardless of the Healthy grade probability. Otherwise, the photos that are not classified as either Referable DR or Non-referable DR, are classified as Healthy. Here, we experimented with the cascading thresholds limits of (∼, 0.4, 0.4) and (∼, 0.3, 0.3), which are formatted for the corresponding classes: Healthy, Non-referable DR and Referable DR.

### Margin Max

To our knowledge, the ‘margin max’ technique has not previously applied in similar studies. In this method, if the top two less sever classes’ probabilities (e.g. Healthy and Non-referable DR) are within a set given threshold, to boost the sensitivity of a certain class (e.g. Healthy) the maximum rule will be ignored. As an example, consider the case of the ‘margin max’ of (0.2) for boosting the sensitivity of the Healthy grade. If the Softmax scores of Healthy, Non-referable and Referable DR were assigned as [0.3 −0.45 −0.25] respectively, the Healthy grade is chosen although it is not the maximum of three probabilities.

### Microsoft Azure Parameters

A standard NV6 Windows instance (6 VCPUs, 56 GB memory) from East US 2 region was selected as the training virtual machine. An additional standard SSD disk of 1023 GB storage space was attached to the training virtual machine [Figure 4].

**Figure 4:**
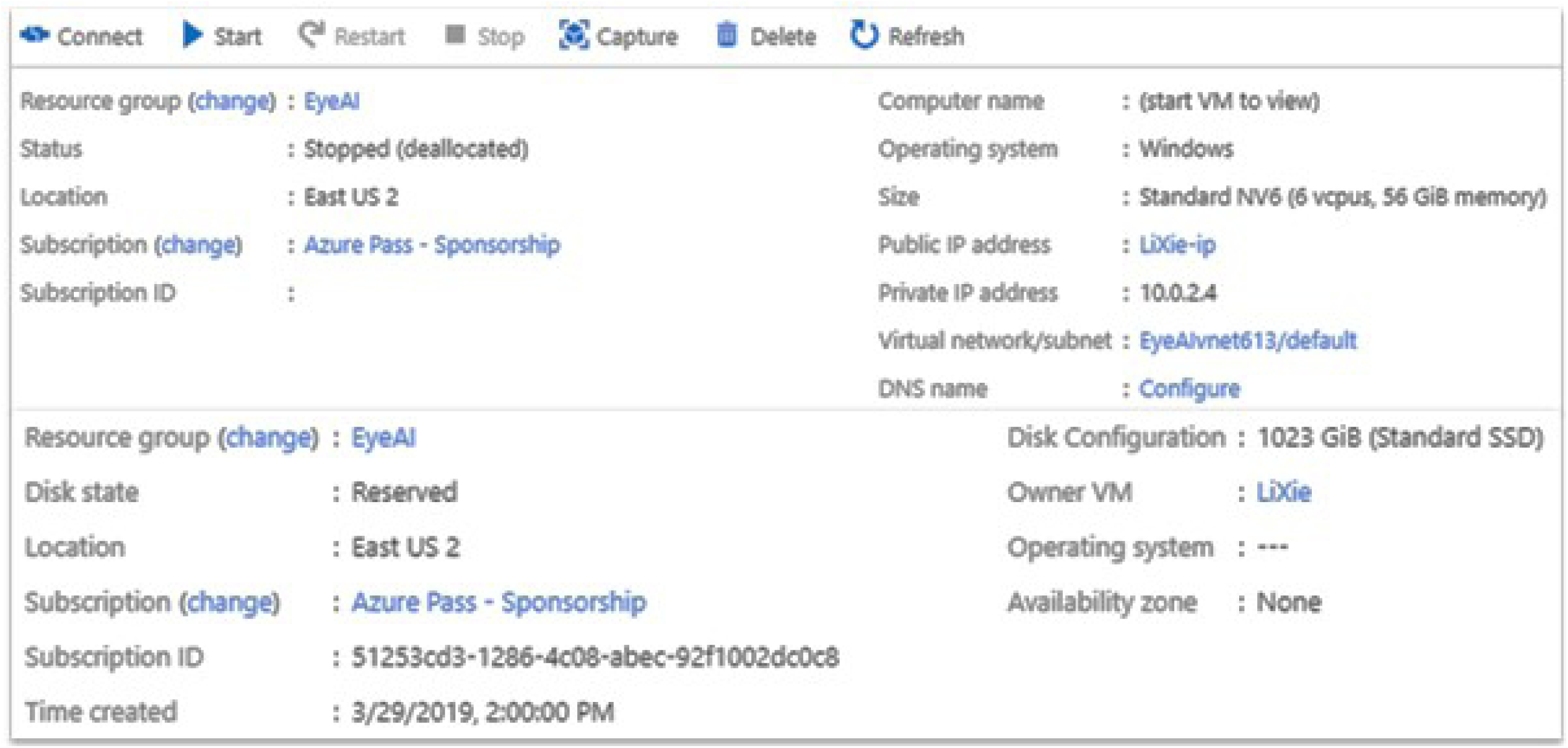
Screenshot of the Microsoft Azure™ virtual machine. A Virtual Machine was created on Microsoft Azure East US server. 6 CPUIs were avaialble to us on this Virtual Machine, and it was used for training and validation process.

## Results

### Generate the curated testing set

The “Quality Assessment” CNN reached 99% accuracy and the validation loss of lower than 0.05. This CNN model was then used to create a ‘curated’ test set from the Kaggle EyePACS dataset. The ‘curated’ test set included 6,900 images from the original 13,305 ‘un-curated’ set (i.e. a 47% rejection rate).

### Un-curated testing set versus curated testing set

The “Classifier” CNN was trained and validated using the Microsoft Azure™ cloud platform. This was done twice, once for the binary DR grading classification Healthy vs Diseased, and once for the tertiary DR grading classification Healthy, Non-referable DR, and Referable DR. The cross-entropy and accuracy were tracked and recorded throughout the training and validation process. The training progress was monitored for 100 epochs and the best set of weights that resulted in minimal validation loss was picked and set for the proceeding CNN performance assessment.

While, the “Classifier” CNN was trained and validated using ‘un-curated’ data, it was tested separately using unseen ‘curated’ and ‘un-curated’ data. One would assume that using ‘curated’ (i.e. higher quality) data for the CNN test would improve the performance of the model. Here and for the first time, we wanted to assess this hypothesis [Table 3&4].

**Table 3:**
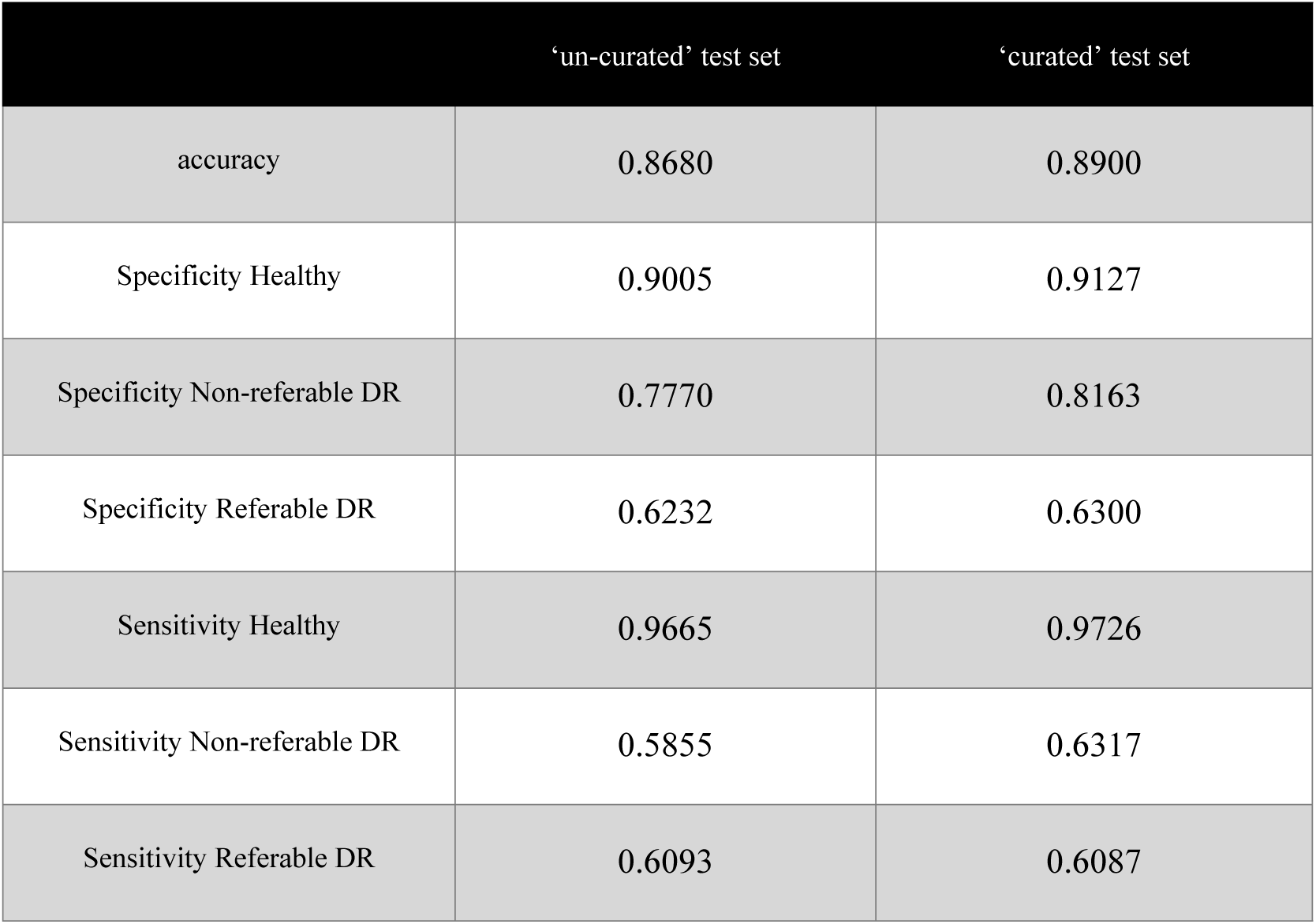
Performance of the ‘classifier’ CNN based on three grades (Healthy, Non-referable, Referable)

|  | ‘un-curated’ test set | ‘curated’ test set |
| --- | --- | --- |
| accuracy | 0.8680 | 0.8900 |
| Specificity Healthy | 0.9005 | 0.9127 |
| Specificity Non-referable DR | 0.7770 | 0.8163 |
| Specificity Referable DR | 0.6232 | 0.6300 |
| Sensitivity Healthy | 0.9665 | 0.9726 |
| Sensitivity Non-referable DR | 0.5855 | 0.6317 |
| Sensitivity Referable DR | 0.6093 | 0.6087 |

**Table 4:** Performance of the ‘classifier’ CNN based on two grades (Healthy, Diseased)

|  | ‘un-curated’ test set | ‘curated’ test set |
| --- | --- | --- |
| accuracy | 0.896 | 0.909 |
| Specificity Healthy | 90.05 | 91.27 |
| Specificity Diseased | 88.01 | 89.43 |
| Sensitivity Healthy | 96.65 | 97.26 |
| Sensitivity Diseased | 69.75 | 71.32 |

Interestingly, the “Classifier” CNN prediction performance improved only marginally for the ‘curated’ test sets, compared to the ‘un-curated’ set.

### Sensitivity Uplift

Several implementations of ‘cascading thresholds’ and ‘margin max’ techniques were then used to boost the sensitivity of the “Classifier” CNN, using the ‘curated’ and ‘un-curated’ test sets, for both two and three grading level schemes.

It appeared that Cascading Thresholds (∼, 0.3, 0.3) and Margin Max (0.4) were the most effective techniques for sensitivity boosting. We then investigated the effects of these techniques to boost the sensitivity of CNN towards either the Healthy or most Diseased grade [Tables 5-8].

**Table 5:** sensitivity boost of the ‘curated’ dataset with three labels, for Healthy and Diseased categories

| | Margin Max<br>(0.4)<br>Boosting<br>Healthy | Origina<br>1 | Cascading Thresholds (∼, 0.3,<br>0.3)<br>Boosting Diseased |
| --- | --- | --- | --- |
| accuracy | 0.8827 | 0.89 | 0.872 |
| Specificity Healthy | 0.8957 | 0.9127 | 0.9291 |
| Specificity Non-referable<br>DR | 0.8374 | 0.8163 | 0.7245 |
| Specificity Referable DR | 0.6954 | 0.63 | 0.5284 |
| Sensitivity Healthy | 0.9852 | 0.9726 | 0.9364 |
| Sensitivity Non-referable<br>DR | 0.5755 | 0.6317 | 0.6676 |
| Sensitivity Referable DR | 0.5072 | 0.6087 | 0.7199 |

**Table 6:** sensitivity boost of the ‘un-curated’ dataset with three labels, for Healthy and Diseased categories

|  | Margin Max (0.4)<br>Boosting Healthy | Original | Margin Max (0.4)<br>Boosting Diseased |
| --- | --- | --- | --- |
| accuracy | 0.8680 | 0.868 | 0.844 |
| Specificity Healthy | 0.8840 | 0.9005 | 0.923 |
| Specificity Non-referable DR | 0.8004 | 0.777 | 0.6707 |
| Specificity Referable DR | 0.7624 | 0.6232 | 0.5052 |
| Sensitivity Healthy | 0.9831 | 0.9665 | 0.9154 |
| Sensitivity Non-referable DR | 0.5503 | 0.5855 | 0.6203 |
| Sensitivity Referable DR | 0.5026 | 0.6093 | 0.7522 |

**Table 7:**
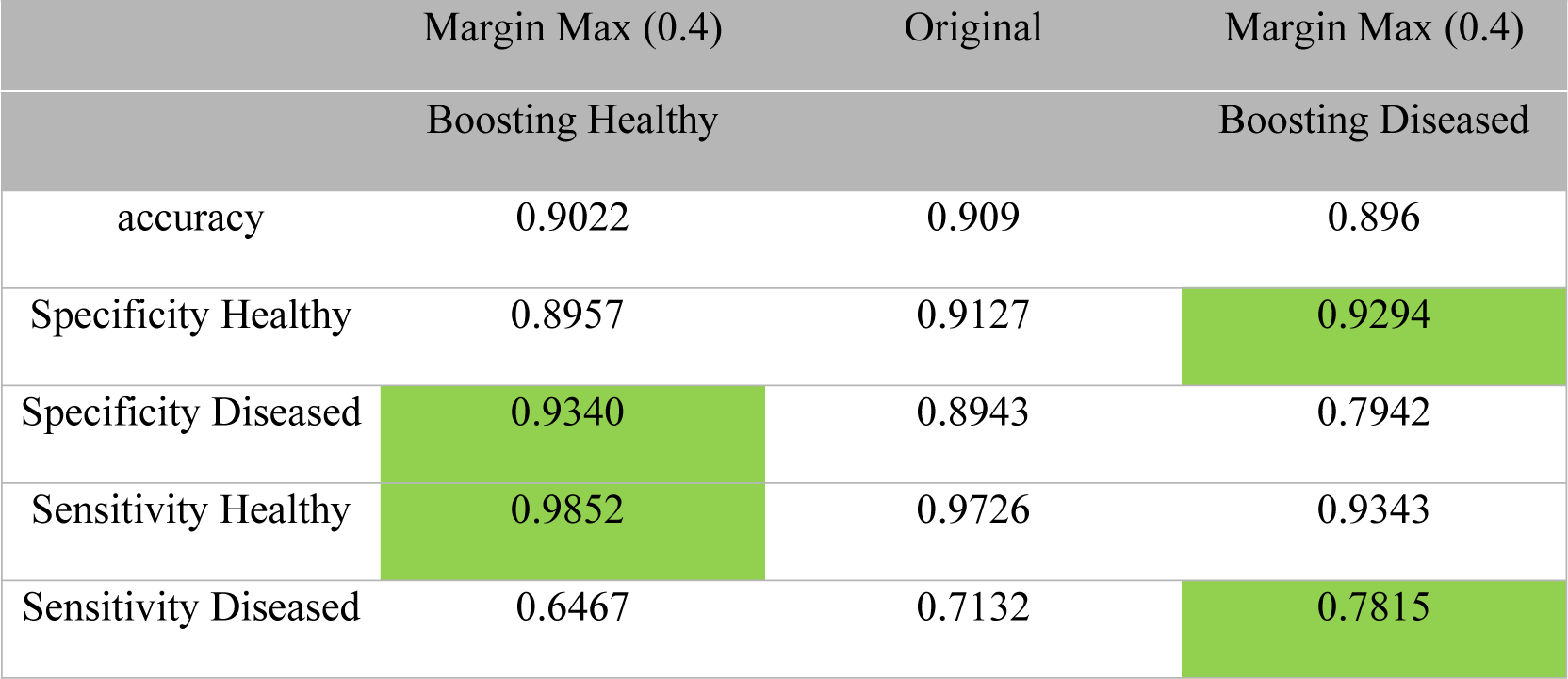
sensitivity boost of the ‘curated’ dataset with two labels, for Healthy and Diseased categories

**Table 8:** sensitivity boost of the ‘un-curated’ dataset with two labels, for Healthy and Diseased categories

|  | Margin Max (0.4) | Original | Margin Max (0.4) |
| --- | --- | --- | --- |
|  | Boosting Healthy |  | Boosting Diseased |
| accuracy | 0.8921 | 0.896 | 0.88 |
| Specificity Healthy | 0.8840 | 0.9005 | 0.923 |
| Specificity Diseased | 0.9297 | 0.8801 | 0.7659 |
| Sensitivity Healthy | 0.9831 | 0.9665 | 0.9154 |
| Sensitivity Diseased | 0.6346 | 0.6975 | 0.7836 |

It appeared that boosting the sensitivity using both ‘cascading thresholds’ and ‘margin max’ had a similar effect for ‘curated’ and ‘un-curated’ datasets. Also, it seemed that uplifting the sensitivity of the Healthy grade, also enhanced the specificity of the Diseased state, and vice versa.

Here we achieved specificity and sensitivity as high as 89% and 98%, using a bi-classification grading scheme. Here we have shown [Table 7 & 8] that by tweaking the post processing of the outcome of a CNN, we have outperformed the previously published best performance of Kaggl EyePACS, which was later failed to be replicated(35).

## Discussion

Diabetic retinopathy is the most common microvascular complication of diabetes and is the leading cause of blindness among the working-age population (42). Whilst the risk of sight loss from DR can be reduced through good glycaemic management (43), if sight-threatening DR develops, timely intervention with laser photocoagulation or injections of anti-vascular endothelial growth factor (44, 45). The risk of sight loss is to be reduced, patients with diabetes should have their eyes screened regularly to facilitate the detection of DR whilst it is still treatable and before vision loss (46). Unfortunately, in many regions including New Zealand, the attendance at DR screening falls below the recommended rates (47-49), and this is particularly true for those who live in remote areas and those of lower socioeconomic status (50-52).

Whilst eye-screening programs have now been established in many Western Health economies, significant challenges exist to ensure that the service is both equitable and all patients at risk are screened regularly. These challenges include the need for a team of trained clinicians to read the photographs, the high capital cost of retinal cameras to take the photographs, and an efficient administrative IT support system to run it. All these challenges are more acute in the developing world which is known to have a general shortage of healthcare professionals (53).

Incorporating AI to accurately grade the fundus images for DR would offer many benefits to DR screening programs; reducing their reliance on trained clinicians to read photographs, enabling point of contact diagnosis reducing the need for complex IT support systems as well as identifying those patients who need a referral to Ophthalmology services on the day of screening.

Research into AI design and its development for DR screening has progressed significantly in recent years, and this field has enjoyed a good deal of attention of late (54-56). However, for all the excitement none of this work has progressed to a clinically useful tool, providing a real-world AI-solution for DR screening programs. This is due largely to the inability of the research-driven AI to generalize to a real-world setup. Whilst there are many reasons for such a lack of generalisation, the principal ones are the use of small and ‘curated’ datasets and an emphasis on overall accuracy, rather than sensitivity of the developed AI. The AI’s reliance on powerful computers that are not available in most clinical environments has been an additional contributory factor.

During this research, we endeavoured to address those issues that hinder the clinical translation of an in-house developed AI for DR screening. Our “Classifier” CNN was developed and tested using real-world ‘un-curated’ data. Here we demonstrated that our “Classifier” CNN is ‘robust’, as its performance is not critically affected by the quality of the input data. Furthermore, this process of data management, model training and validation was performed using Microsoft’s Azure™ cloud platform. In doing so, we have demonstrated that one can build AI that is constantly re-trainable and scalable through cloud computing platforms. Although few DR AIs are accessible online, to our knowledge this is the first time that an AI is fully implemented and re-trainable through a cloud platform. Hence, provided there is internet access, our AI is capable of reaching remote and rural places; areas traditionally not well served by existing DR screening services.

We have also successfully experimented with two “sensitivity-boosting” techniques, ‘cascading thresholds’ and the ‘margin max’ technique. We observed good improvements in sensitivities and specificities of either Healthy or Diseased grades, depending on the application mode. In doing so we boosted the AI’s sensitivity to detect Healthy cases to more than 98%, (while also improving the specificities of the other more severe classes). These techniques also boosted the AI’s sensitivity of referable disease classes to near 80%.

The sensitivity of a screening test is the percentage of the condition that is correctly detected; the specificity of a screening test is the percentage of people that one refers unnecessarily. Within all screening programs, the need to balance high sensitivity with an acceptable specificity has been long recognised. Traditional diabetic eye screening programs, therefore mandate a minimum sensitivity of >85% and specificity of >80% for detecting sight-threatening diabetic retinopathy as there is a personal and financial cost associated with unnecessary referrals to eye clinics (57). Although we have not chosen categories of non-sight-threatening DR and sight-threatening DR in this paper, clearly the CNN classifier we report here would not achieve these stringent targets. However, before rejecting the performance of the CNN we need to consider the role that a classifier CNN could play in a diabetic eye screening program. Whilst it is appropriate that a screening program has to strike the correct balance between both a high sensitivity and specificity, we envisage that in many situations, a classifier CNN will not be the sole arbitrator for grading diabetic retinopathy.

If one then considers the CNN simply as an adjunct to the wider program, using the techniques we describe in this paper the opportunity to develop classifier CNN’s that are tailored to the specific requirements of the program becomes possible (58). The United Kingdom national Ophthalmology database study; revealed that Of the 48,450 eyes with structured assessment data at the time of their last record, 11, 356 (23.8%) eyes had no DR (59). Thus a sensitivity boosted classifier like the one described here, manipulated to detect healthy eyes with high sensitivity could be used to rapidly and safely triage eyes with DR from eyes with no DR (healthy eyes) reducing the number of images sent for structured grading by over 20%. Although such an approach may have appear to having limited utility in the context of traditional screening, a CNN which removes the need for a significant percentage of images to be sent for human review, would lead to immediate and significant cost savings for the program.

In the context of a rural setting, the ability to identify those patients with no disease from those with any disease is particularly valuable. Whilst the aim of diabetic eye screening programs thus far has been to detect sight-threatening DR it is well recognised that patients with even mild DR are at increased risk of progression compared with no DR, and the rate of progression increases with the level of DR (60). The development of any DR is therefore a significant event and one that could be used to target valuable and scarce health care resources more effectively to those at highest risk. In effect, even if no other CNN approach was then used, the relatively simple cloud-based CNN we describe here would help identify those patients at increased risk of either advanced disease or disease progression, and who therefore merit further review. Moreover, using the techniques described here, more sophisticated classifier CNNs could also be developed, ones that are manipulated to detect disease (with a very high sensitivity). It is conceivable that different classifier CNNs could then be run concurrently within diabetic eye screening programs to sequentially grade differing levels of disease with high sensitivity, ultimately leaving the human grading team with a relatively small number of images to review for adjudication and quality assurance.

Arguably, one of the biggest challenges that faces all AI-based “diagnostic” systems is the issue of public trust. Whereas it is accepted that in a screening program with a sensitivity of 90%, 1 in 10 patients will be informed that are healthy when in actual fact they have diseases, well-publicised failures of AI systems suggest that the public would not accept such failure rates from a “computer” (61). Whilst the relatively simple CNN described in this paper lacks the required sensitivity to be the ***sole*** arbitrator for identifying ***referable*** disease in a structured screening program, the fact that the methods we describe boosted the sensitivity of the CNN to detect disease by over 10% in most cases is noteworthy. We therefore believe that the techniques we describe here will prove to be valuable tools for those looking to build bespoke CNN’s in the future.

In conclusion, we have demonstrated how existing machine learning techniques can be used to boost the sensitivity of a CNN classifier to detect both health and disease. We have also demonstrated how even a relatively simple classifier CNN, one that is capable of running on a cloud-based provider, can be utilised to support both existing DR screening programs and the development of new programs serving rural and hard to reach communities. Further work is required to both develop classifiers that can detect sight-threatening DR with a very high sensitivity, and evaluate how a battery of CNN’s each with differing specifications and roles, may be used concurrently to develop a real-world capable, fully automated DR screening program.

## Acknowledgements

This study was made possible by a Microsoft Azure Asia – Cloud Research Fellowship

## Author contributions

Dr Ehsan Vaghefi performed the full analysis, and wrote bulk of the manuscript including the Methods and Results

Mr Song performed part of AI design and testing. He also contributed to the first draft of the paper.

Dr Xie performed part of AI design and testing. He also contributed to the first draft of the paper.

Dr David Squirrell supervised the entire project and contributed to the introduction and discussion sections.

## Data availability

We have used publically available dataset, which can be found here: https://www.kaggle.com/c/diabetic-retinopathy-detection/data.

